# An electrostatic-interaction-based mechanism triggering misfolding of prion proteins from cellular isoform to scrapie isoform

**DOI:** 10.1101/851493

**Authors:** J. Li, X. Ma, S. Guo, C. Hou, L. Shi, L. Ye, L. Yang, B. Zheng, X. He

## Abstract

Understanding how prion proteins refold from a cellular isoform (PrPC) to a disease-causing isoform (PrPSc) has been among the “ultimate challenges” in molecular biology, biophysics, pathology, and immunology. Conformational changes of prion proteins from PrPC to PrPSc involve the unfolding of a short α-helix that overshadows the challenge. Considering the mechanisms of electrostatic attraction, thermal disturbance, hydrogen ion concentration, hydrophobic interaction, and the shielding effect of water molecules, this study reveals an electrostatic-interaction-based mechanism by means of which prion proteins refold in an aqueous environment. The electrostatic-interaction-induced protein unfolding mechanism causes a hydrophobic polypeptide segment to dangle out over the conglobate main body of the prion protein, thereby allowing the first triangular hydrophobic rung formation via hydrophobic interaction. A molecular model of PrPSc is proposed that allows the β-solenoid with a triangular hydrophobic core.

**Statement of Significance:** We present three main results that would revolutionize the understanding of pathology of prion diseases. First, the prion protein refolding (from cellular isoform to scrapie isoform) derives from the unfolding of the shortest α-helix of PrPC, which provides a long polypeptide segment full of hydrophobic residues dangling out over the conglobate main body of the prion protein, thereby allowing formation of the first triangular hydrophobic rung via hydrophobic interaction. Second, polyanions-induced increasing in local concentration of hydrogen ion (i.e., the PH increase) undermines the shielding effect of water molecules, thereby allowing escape of the arginine side chains from the hydration shell, destabilizing the shortest α-helix and initiating the refolding of PrPC. Third, a β-solenoid structural model for PrPSc with a triangular hydrophobic core is proposed.

## Introduction

Prion diseases are several fatal neurodegenerative diseases caused by misfolded prion proteins in humans and many other mammals(*1*). In humans, prions are believed to be the cause of Creutzfeldt–Jakob disease, fatal familial insomnia, kuru, and Gerstmann–Sträussler–Scheinker syndrome(*2*). There are two distinct isoforms of prion protein: the normal isoform of the protein is called the cellular prion protein (PrP^C^) and the infectious isoform is called the scrapie prion protein (PrP^Sc^). Currently, the generally accepted theory in the field is that a molecular structural change in prion proteins (from PrP^C^ to PrP^Sc^) is the cause of prion diseases and their infectiousness. The molecular pathogenesis of prion diseases is expressed by a misfolded three-dimensional (3D) structure of the PrP^Sc^ that derives from the refolding of the PrP^C^(*3*). Since Prusiner won the 1997 Nobel Prize in Physiology or Medicine for his work revealing the new biological principle of infection of prions (*4*), understanding how prion protein refolds from the cellular isoform to the scrapie isoform has become a fundamental task in molecular biology, structural biology, biological physics, pathology, and immunology(*5*). In the past decade, experimental evidence has suggested that many other age-related neurodegenerative diseases (such as amyotrophic lateral sclerosis, Alzheimer’s disease, Parkinson’s disease, and Huntington’s disease) have pathogenic mechanisms similar to prion diseases, and these diseases have been termed “prion-like” diseases (*6, 7*). This means that the protein refolding mechanism of prions may play a part in the process of these age-related neurodegenerative diseases. Hence, understanding the mechanism of prion refolding (from PrP^C^ to PrP^Sc^) has broad pathological significance and is critical for understanding prion diseases and for developing anti-prion therapeutics(*8*). However, despite today’s most advanced experimental techniques, it is not known what causes the conversion of the normal form to the infectious form of prions.

A PrP^C^ molecule consists of a partially unstructured N-terminal domain (residues 23–125), and a globular C-terminal domain (residues 126–230). The globular C-terminal domain comprises three α-helices of different lengths (α-helix content 30%) and almost no β-sheet (*9*)(see Fig. 1a). Thus, PrP^C^ can be considered a partially unfolded protein without well-defined molecular structure in the N-terminal domain(*10*). Although the exact 3D structure of PrP^Sc^ is not known, Fourier transform infrared (FTIR) spectroscopy evidence has shown that a large β-solenoid structure (β-sheet content about 43%) appears in place of the shortest α-helix structure after the transformation from PrP^C^ to PrP^Sc^, causing the well-defined secondary structure content increase from 30% to 65% (*3*). This finding means that the transformation from PrP^C^ to PrP^Sc^ can be conceived as a partial refolding process of the protein. However, it is surprising that the shortest α helix of PrP^C^ is found to be unfolded during PrP^Sc^ formation while the other two α-helices are well preserved after PrP^Sc^ formation (see Fig. 1a), because the structural stability of the α-helix is much greater than that of the other secondary structures(*11-13*). No experimental evidence suggests that the phenomenon of α-helix-refolding exists widely in nature. It most likely that an undetected force or mechanism unfolds the shortest α-helix, allowing a structured polypeptide segment to fall into an unfolded state during the protein refolding process. Otherwise it is difficult to explain how the well-structured α-helix can refold into β-solenoid architecture without the unfolding process during PrP^Sc^ formation. Unfolding of the shortest α-helix is seen as a necessary condition for activating the transformation from PrP^C^ to PrP^Sc^.

**Figure 1.**
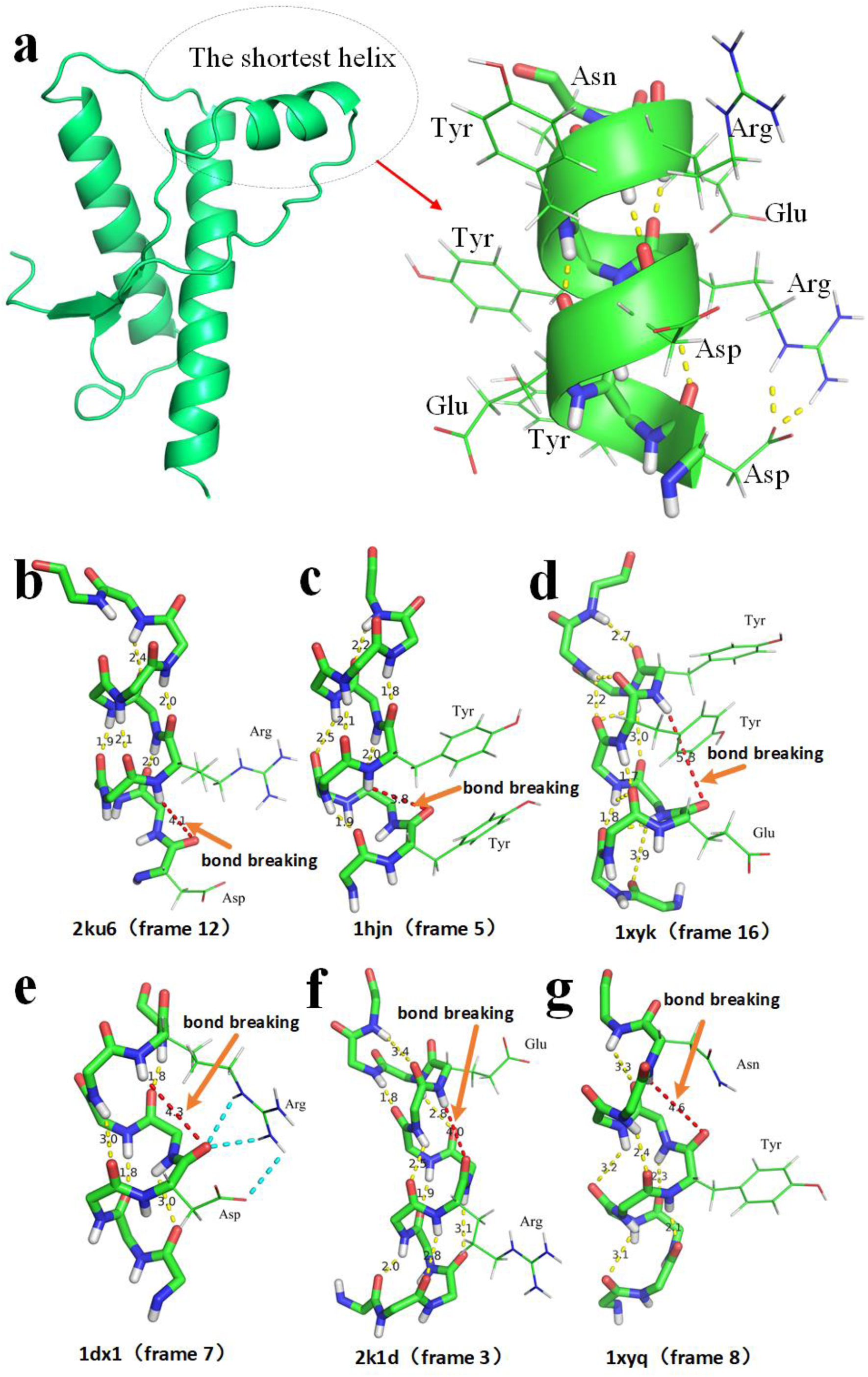
Intrinsic structural instability of the shortest α-helix of PrP^C^, providing experimental evidence that breakage of all hydrogen bonds in the α-helix can be found in PDB. **a**, The globular C-terminal domain consists of three α-helices of different length. **b**, the shortest α-helix of 2ku6. **c**, the shortest α-helix of ihjn. **d**, the shortest α-helix of 1xyk. **e**, the shortest α-helix of 1xy1. **f**, the shortest α-helix of 2k1d. **g**, the shortest α-helix of 1xyq.

## Discussion

Anfinsen’s dogma claimed that the native structure of a small protein is completely determined by the protein’s amino acid sequence(*14*). Therefore, there should be some physical folding codes in the amino acid sequence that dictate the formation of β-solenoid architecture after the unfolding process. These physical folding codes are most likely activated by thermal-motion-induced hydrophobic interactions forming the β-solenoid architecture that underlies the transformation from PrP^C^ of PrP^Sc^(*15*). The physical folding codes for the β-solenoid architecture formation may have low priority; otherwise it is difficult to explain why these folding codes cannot be activated during the protein folding process of PrP^C^. In any case, unfolding of the shortest α-helix provides the precondition for formation of the β-solenoid architecture. An α-helix possesses outstanding structural stability, due to maximization of the number of formations of intramolecular hydrogen bonds of the main chain (*11, 12, 16, 17*). Structural instability of the shortest α helix of PrP^C^ in cellular environments should be regarded as a considerable anomaly or mystery. Thus, in this problem lies our lack of understanding of the reason why the shortest α-helix of PrP^C^ can unfold during the transformation from PrP^C^ to PrP^Sc^. The mechanism by which water molecules interact with the shortest α-helix of PrP^C^ may explain the one of the great mysteries of prion diseases. In general, organic solvents denature proteins(*18*). That means that water molecules are most likely indispensable to structural instability in proteins. Hydrophilic side chains of proteins are normally hydrogen bonded with surrounding water molecules in aqueous environments, thereby preventing the side chains from interfering with the stability of the secondary structure(*19*).

Many native structures of PrP^C^ of different species have been experimentally determined using a solution nuclear magnetic resonance (NMR) method that provides experimental evidence for the intrinsic structural instability of the shortest α helix of PrP^C^ in aqueous environments. Many experimental results have shown breakage of the intramolecular hydrogen bonds of the shortest α-helix of PrP^C^, demonstrating the instability of the α-helix in aqueous environments10.20-24(*10, 20-24*), (see Fig. 1b∼g and Supplementary S2). From analysis of 200 protein structures in the Protein Data Bank (PDB) archive, it can be found that the distance between a carbonyl oxygen atom and its hydrogen bonded (paired) amide hydrogen atom in an α-helix is about 2.3 Å (see Supplementary S1). However, in these experimentally determined shortest α-helix structures of PrP^C^s, the distances between some paired neighboring C=O and N-H groups have been measured as clearly greater than 3.4Å, causing the interaction between the paired C=O and N-H groups to degrade from hydrogen bonds to weak electrostatic attraction (see Fig. 1b∼1g). Note that the shortest α-helices in different PrP^C^s always consist of polypeptide segments of a same amino acid sequence (EDRYYRE), from which we can conceive that these α-helices are the same (see Supplementary S3). Moreover, experimental evidence for the breakage of all the hydrogen bonds of an α-helix can be found, as shown in Fig. 1 and Supplementary S2. From analysis of the protein structures in PDB, we find that the structural instability of the shortest α helix at least prevailed in 16 different prion proteins (see Supplementary S2). All these unstable native structures of prion proteins were experimentally determined using a solution-NMR method. These findings indicate that the intrinsic instability of the shortest α-helix of PrP^C^ may be resourced from interaction between the α-helix and surrounding water molecules.

If, on the basis of experimental evidence, we can clearly explain why the shortest α-helix demonstrates intrinsic structural instability (i.e., breakage of intramolecular hydrogen bonds of the main chain) in aqueous environments, we may be able to reveal that transformation from PrP^C^ to PrP^Sc^ is triggered by unfolding of the shortest α-helix as a key pathogenesis mechanism of prion diseases. Aspartic acid (D) and glutamate (E) contain negatively charged side chains. Arginine (R) and lysine (K) contain positively charged side chains. Tyrosine (Y) contains a flattened hydrophobic side chain. In mammalian prion proteins, the fixed amino acid sequence (EDRYYRE) threads the shortest α-helix of PrP^C^. Experimental evidence suggests that the breakage of two hydrogen bonds of the α-helix is caused by the formation of hydrogen bonds between amide hydrogen atoms of the R residues and carbonyl oxygen atoms of the main chain (see Figs. 1e, S1i, and S1j). This phenomenon indicates that the arginine side chain is capable of interfering with the stability of the secondary structure through the formation of hydrogen bonds with the main chain (see Figs. 1e, S1i, and S1j). In accordance with the CHARMM force field, the amide hydrogen atom of R residue in molecular models has a charge of 0.46e, which is obviously greater than the charge (0.31e) of the amide hydrogen atom in the main chain of proteins (see Fig. 2). Therefore, negatively charged carbonyl oxygen atoms of the main chain would be more likely to hydrogen bond with the side chain of the R residue than would an amide hydrogen atom of the main chain (see Figs. 1e, S1i, and S1j).

**Figure 2.**
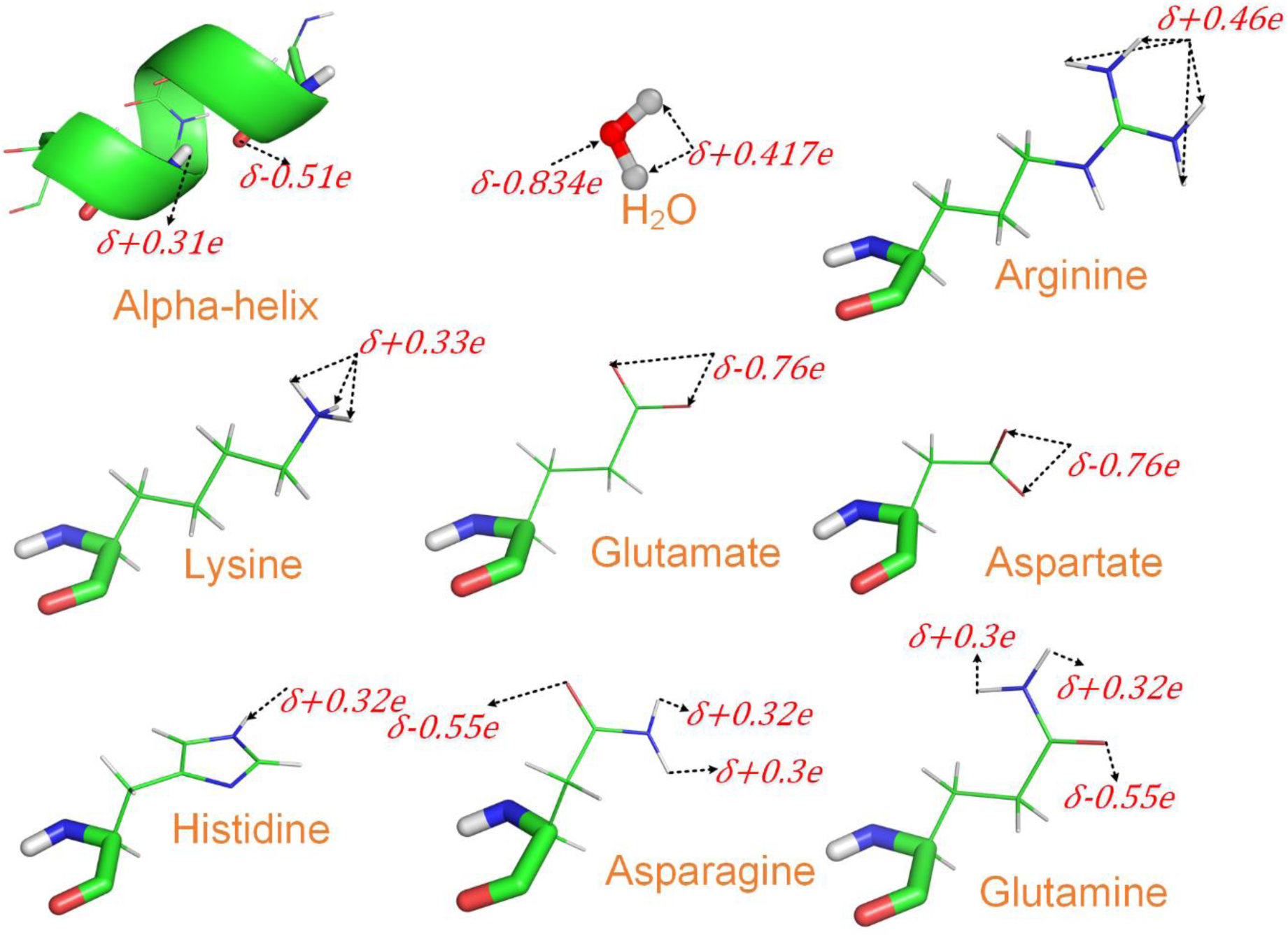
Comparison of charges among water molecules, charged side chains, carbonyl oxygen atoms, and amide hydrogen atoms of the main chains.

## Method

To further demonstrate the capability of the arginine side chain to interfere with the stability of the α-helix, we tried to minimize the potential energy of the shortest α-helix in isolation, using a CHARMM force field(*25*) and a microcanonical (NVE) ensemble relaxation engine (NVERE)(*26*). The simulation results showed that the potential energy of the α-helix could be further minimized in isolation through the formation of hydrogen bonds between side chains of the R residues and carbonyl oxygen atoms of the main chain (see Fig. 3). It is worth noting that hydrophilic side chains are normally hydrogen bonding with hydration shells in an aqueous environment, due to the strong polarity of water molecules (see Fig. 2). This characteristic indicates that surrounding water molecules shield a hydrophilic side chain from hydrogen bonding with C=O groups and N-H groups of the main chain, thereby preventing the side chains from interfering with the formation and stability of secondary structures(*19*). It also explains why secondary structures (such as α-helices and β-sheets) are always stabilized by hydrogen bonds between the N-H groups and C=O groups of the main chain(*19*).

**Figure 3.**
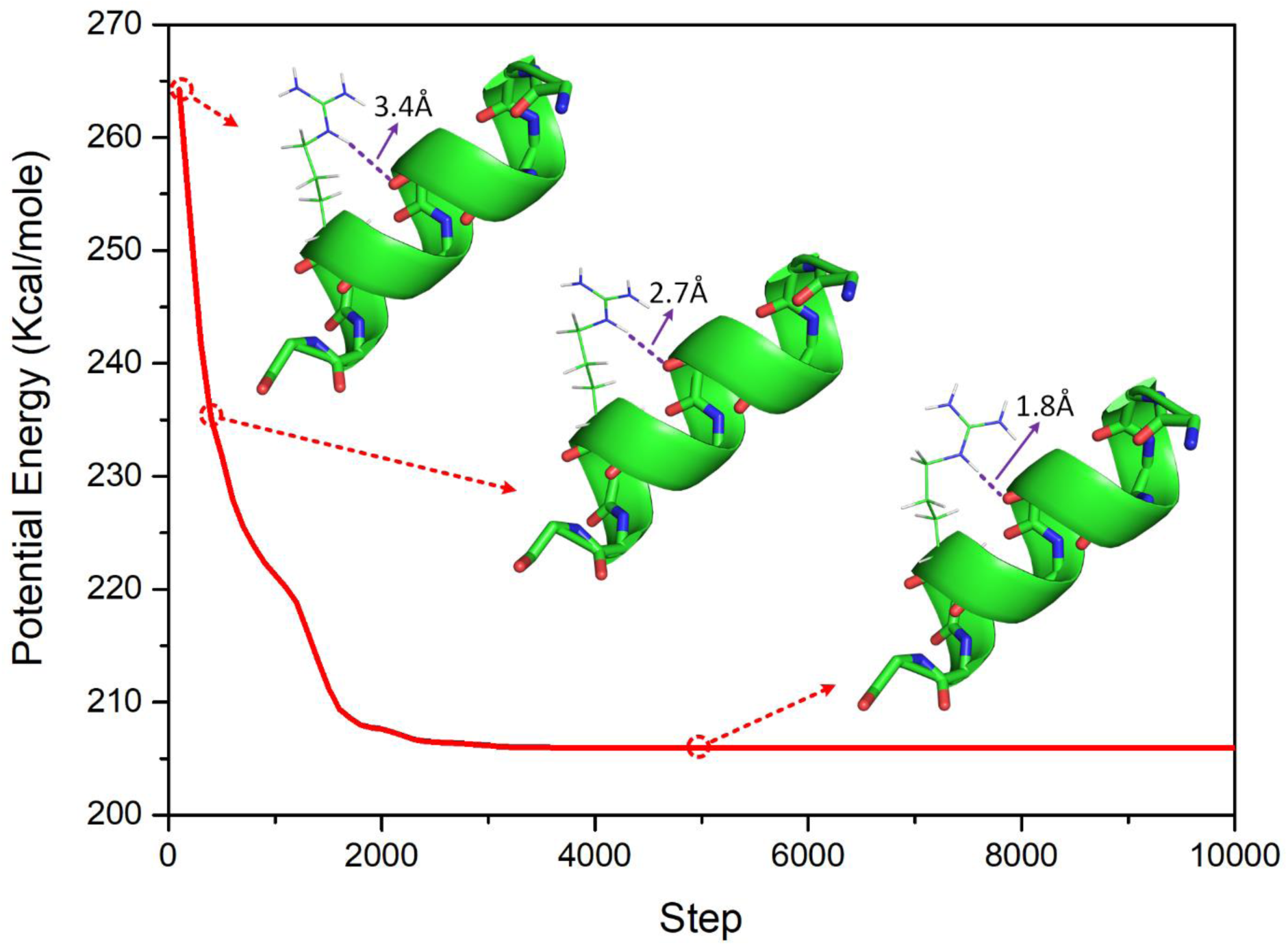
The potential energy of the α-helix could be further minimized in isolation through the formation of hydrogen bonds between side chains of the R residues and carbonyl oxygen atoms of the main chain.

How can the arginine side chain of the α-helix escape from the shielding effect of surrounding water molecules that destabilize the α-helix through hydrogen bonding with the carbonyl oxygen atom of the main chain of the prion proteins? Note that the two arginine side chains are adjacent to the aspartic acid side chain and the glutamate side chain in the EDRYYRE helix. Thermal motion triggers some electrostatic attraction between these charged adjacent side chains, thereby pulling the side chains of arginine close to the main chain and causing hydrogen bonding between the side chains and the carbonyl oxygen of the main chain (see Figs. 1e and S1j). When the hydrophilic group of the arginine residue approaches carbonyl oxygen of the main chain, two hydrogen bonds can be formed between them, as shown in Fig. 1e, causing the carbonyl oxygen atom to escape from the hydrogen bonding with the amide hydrogen atom in the main chain, because each carbonyl oxygen atom allows the formation of only two hydrogen bonds, as shown in Fig. 1e. Experimental evidence also show that the two tyrosine side chains in the α helix also can contribute to the structural instability of the secondary structure by cutting off a hydrogen bond of the main chain due to hydrophobic interactions between them (*27, 28*), as shown in Fig. 1d. However, there must be some other mechanism that contributes to the unfolding of the shortest α-helix of PrP^C^ during the formation of PrP^Sc^, otherwise it would be difficult to explain how PrP^C^s could exist stably in aqueous environments.

It is worth noting that human and murine PrP^C^ undergo a pH-dependent conformational change in the region of pH 4.4–6, with a loss of helix and gain of β-solenoid structures. Thus, low pH most likely plays a role in unfolding the α-helix that ultimately results in PrP^Sc^ formation. Moreover, much of the experimental evidence for the intrinsic instability of the shortest α-helix of PrP^C^ has been achieved in acidic conditions with pH 4.5(*10, 21-23*), as shown in Fig. 1. However, acidic conditions with pH 4.4–6 do exist in the bulk aqueous phase of cytoplasm in nature. Over the last decade, the theory has become more widely accepted that a nucleic acid, such as an RNA molecule, might be a cellular cofactor for PrP^Sc^ formation; such a nucleic acid would act as a catalyst rather than encoding genetic information in that case(*29, 30*).Both RNA and DNA molecules can be classified as polyanions. In solution, polyanions such as DNA and RNA can attract surrounding hydrogen ions and raise the local hydrogen ion concentration. Thus, polyanions such as DNA and RNA in cytoplasm can most likely create a locally acidic environment in cytoplasm. When a polyanion nears a PrP^C^ in cytoplasm, the polyanion is capable of providing an acidic condition to trigger the formation of PrP^Sc^ (see Fig. 4).

**Figure 4.**
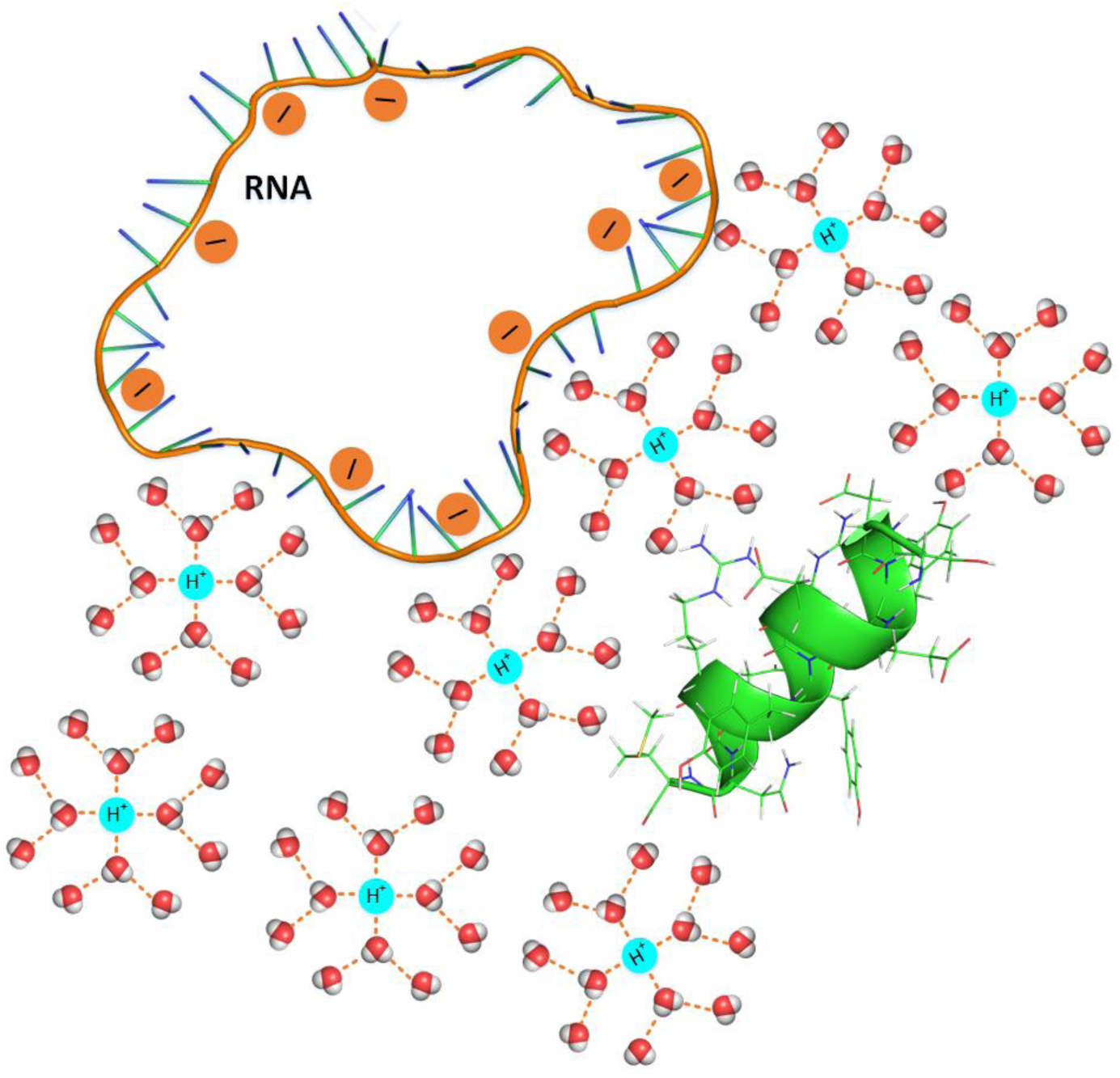
A polyanion-induced increase in the local concentration of hydrogen ions (i.e., a PH increase) undermines the shielding effect of water molecules.

Water molecules are very small and have very strong polarity, allowing them to hydrogen bond with hydrogen bond acceptor groups and hydrogen bond donor groups of each hydrophilic side chain. However, with the increase in hydrogen ion concentration, more and more water molecules would be distributed surround hydrogen ions orderly, due to the hydration shell requirement, releasing the hydrophilic side chain from the shielding effect of water molecules (see Fig. 4). This explain why the arginine side chain of the α-helix is capable of escaping from the shielding effect of surrounding water molecules in the region of pH 4.4–6, allowing hydrogen bonding between the side chain and the carbonyl oxygen atom of the main chain.

From analysis of protein structures in PDB, it can be found that the amino acid sequence of EDRYYRE occurs only in mammalian PrP^C^s (see Supplementary S3). This manifestation indicates that the α-helix-unfolding-induced protein refolding mechanism can exist only in mammalian prion diseases and may explain why prion diseases do not occur in non-mammals.

The molecular structures of two infectious variants, N-terminally truncated PrP^Sc^ (PrP 27-30) and a miniprion (PrP^Sc^106) of the prion protein have been characterized using electron crystallography(*31*). Increasing experimental evidence shows that the unfolded segments of PrP^C^ most likely fold into β-solenoid structures with stacks of triangular rungs and with a hydrophobic core. It is worth noting that only short hydrophobic or neutral side chains can be located within the triangular hydrophobic core (*32*). From analysis of other left-handed and right-handed β-solenoid structures in PDB, lysine, glutamine, glutamate, histidine, asparagine, and aspartate are classified as R_in_ due to their short hydrophobic side chains that can be located within the triangular hydrophobic core. Alanine, arginine, valine, leucine, isoleucine, phenylalanine, tryptophan, and methionine are classified as R_out_ due to their long hydrophilic side chains that cannot occur within the triangular hydrophobic core.

β-solenoid structures consist of β-strands connected laterally by backbone hydrogen bonds(*33*). β-strands are characterized by the parallel distribution of adjacent peptide planes. Such parallel distribution also causes adjacent side chains to be distributed on opposite sides of the main chain, with each side chain parallel to every other side chain. When two R_out_ are linked together in the chain of β-solenoid structures, R_out_-R_out_ segments can only be located at the turns.

Experimental evidence suggests that the β-solenoid of PrP^Sc^ has a triangular hydrophobic core. Note that the unfolding of the shortest α helix brings a long hydrophobic segment of the sequence (AGAAAAGAVVGGLGG) into unfolding status. The first triangular rung may be formed from the hydrophobic segment due to the hydrophobic interaction. In fact, experimental results have shown that G is often present in the turns of proteins’ native structures(*34*). Thus, we can suppose that the G is located at the turns of the first triangular rung. From analysis of amino acid sequences of mammal prions, we propose a β-solenoid structure for PrP^Sc^ with an isosceles triangular hydrophobic core (see Fig.5). The proposed β-solenoid structure is also a good fit for amino acid sequences of many other known prion proteins (see Supplementary S5). Our Fourier transform infrared (FTIR) spectroscopy demonstrated that the β-sheet content of PrP^Sc^ was 43%. Thus, about 99 residues of a prion protein fold into a β-helical structure after the formation of PrP^Sc^. This means that peptides of PHGGGWGQ repeats may be also included in the β-solenoid of PrP^Sc^.

**Figure 5.**
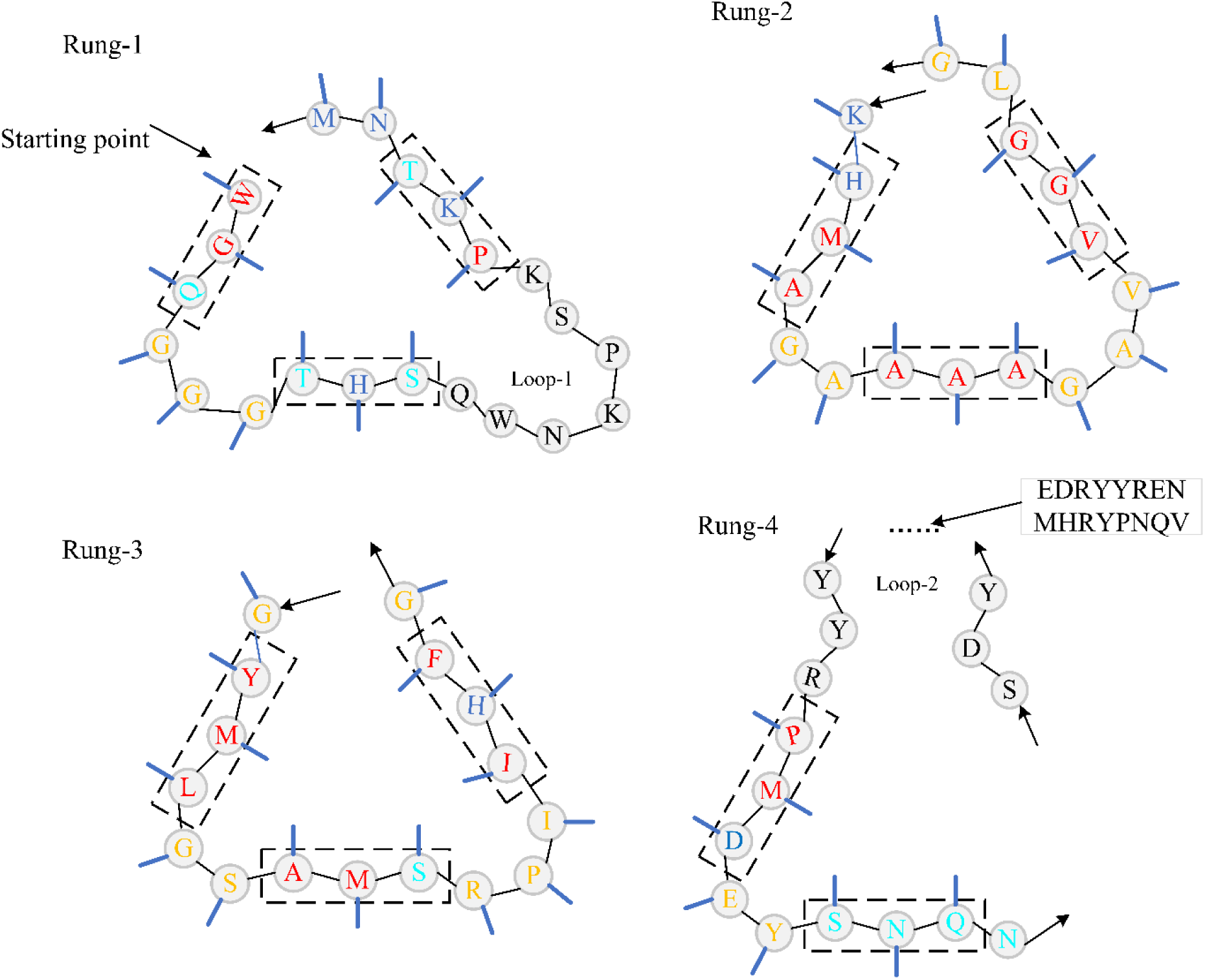
Threading of PrP residues 89–175 onto a left-handed β-helical fold. Inserted loops are indicated.

## Conclusion

In conclusion, the apparent contradiction between the outstanding stability of α-helixes and the fact that β-solenoid is in the place of the shortest α-helix structure after PrP^Sc^ formation should be considered a paradox. The prion protein refolding (from cellular isoform to scrapie isoform) most likely derives from the unfolding of the shortest α-helix of PrP^C^, which provides a long polypeptide segment full of hydrophobic residues dangling out over the conglobate main body of the prion protein, thereby allowing formation of the first triangular hydrophobic rung via hydrophobic interaction. The polyanion-induced increase in the local concentration of hydrogen ions (i.e., the pH increase) most likely undermines the shielding effect of water molecules, thereby allowing escape of the arginine side chains from the hydration shell, destabilizing the shortest α-helix and initiating the refolding of PrP^C^(*21*).Experimental and MD simulation results suggest that thermal vibration triggers some electrostatic attraction between adjacent charged side chains, thereby pulling the arginine side chain out of the hydration shell and creating hydrogen bonding between the arginine side chain residue and the carbonyl oxygen of the main chain. Polyanion-induced concentration of hydrogen ions and electrostatic attraction among neighboring charged side chains work cooperatively to unfold the shortest α-helix, enabling the refolding of the β-solenoid. A β-solenoid structural model for PrP^Sc^ with a triangular hydrophobic core is proposed. The β-solenoid structure model is verified by comparing the results available from experiments in the literature.

## ACKNOWLEDGEMENTS

The authors acknowledge financial support from the Fundamental Research Funds for the Central Universities of China. Lin Yang is indebted to Daniel Wagner from the Weizmann Institute of Science and Liyong Tong from the University of Sydney for their support and guidance. Lin Yang is grateful for his research experience in the Weizmann Institute of Science for inspiration. The authors acknowledge the financial support from the National Natural Science Foundation of China (Grant 21601054), Shenzhen Science and Technology Program and the University Nursing Program for Young Scholars with Creative Talents in Heilongjiang Province of China (Grants UNPYSCT-2017126). The authors thank Joan Rosenthal for editing this paper.

## Additional Information

The authors declare no competing financial interests.

## Author Contributions

L.Yang, L.Ye and X.H. formulated the study. J.L., X.M., S.G., L.Yang, and C.H. conducted the MD simulation. L.Yang, L.Ye and X.H. wrote the paper, and all authors contributed to revising it. All authors discussed the results and theoretical interpretations.

